# Gene model for the ortholog of *DENR* in *Drosophila simulans*

**DOI:** 10.1101/2025.10.13.682112

**Authors:** Megan E. Lawson, Kylee Sanow, Brita Lundeen, Noel Kathryn Bynum, Scott Tanner, Chinmay P. Rele, Kellie S. O’Rourke

## Abstract

Gene model for the ortholog of *Density regulated protein* (*DENR*) in the May 2017 (Princeton ASM75419v2/DsimGB2) Genome Assembly (GenBank Accession: GCA_000754195.3) of *Drosophila simulans*. This ortholog was characterized as part of a developing dataset to study the evolution of the Insulin/insulin-like growth factor signaling pathway (IIS) across the genus *Drosophila* using the Genomics Education Partnership gene annotation protocol for Course-based Undergraduate Research Experiences.

## Introduction

The insulin signaling pathway is a highly conserved pathway in animals and is central to nutrient uptake (Hietakangas & Cohen 2009; Grewal 2009). Density regulated protein was first discovered in a human teratocarcinoma cell line because its concentration in cells increased with cell density (Deyo et al. 1998). Subsequent bioinformatic and biochemical analyses showed that the protein is conserved across eukaryotes and functions in non-canonical translation initiation (Fleischer et al. 2006; Skabkin et al. 2010). *D. melanogaster* flies homozygous for a null, knockout allele of the gene encoding Density regulated protein, *DENR* (FBgn0030802), die as pharate adults, showing a larval-like epidermis and reduced proliferation of histoblast cells (Schleich et al., 2014). Subsequent experiments using both RNAi in S2 cells and the knockout allele in larvae showed that DENR is required, along with its interacting partner MCT-1, for the proper expression regulation of a subset of transcripts required for cell cycle progression and growth. In particular, the loss of *DENR* reduces expression of the insulin receptor and makes larvae less sensitive to insulin signaling (Schleich et al., 2014), thus implicating DENR in the regulation of the insulin signaling pathway (Laskowski et al. 2023).

We propose a gene model for the *D. simulans* ortholog of the *D. melanogaster* Density regulated protein (*DENR*) gene. The genomic region of the ortholog corresponds to the uncharacterized protein LOC6726200 (RefSeq accession XP_039152449.1) in the May 2017 (Princeton ASM75419v2/DsimGB2) Genome Assembly of *D. simulans* (GenBank Accession: GCA_000754195.3). This model is based on RNA-Seq data from *D. simulans* (Graveley et al. 2011; SRP006203) and *DENR* in *D. melanogaster* using FlyBase release FB2023_02 (GCA_000001215.4; Larkin et al., 2021). *D. simulans* is part of the *melanogaster* species group within the subgenus *Sophophora* of the genus *Drosophila* (Sturtevant 1939; Bock & Wheeler 1972). It was first described by Sturtevant (1919). *D. simulans* is a sibling species to *D. melanogaster*, thus extensively studied in the context of speciation genetics and evolutionary ecology (Powell 1997). Historically, *D. simulans* was a tropical species native to sub-Saharan Africa (Lemeunier et al. 1986) where figs served as a primary host (Lachaise & Tsacas 1983). However, *D. simulans’s* range has expanded world-wide within the last century as a human commensal using a broad range of rotting fruits as breeding sites (https://www.taxodros.uzh.ch; Lawson et al. 2024). The Genomics Education Partnership maintains a mirror of the UCSC Genome Browser (Kent WJ et al., 2002; Gonzalez et al., 2021), which is available at https://gander.wustl.edu.

## Results

### Synteny

The target gene, *DENR*, occurs on chromosome X in *D. melanogaster* and is flanked upstream by *CG4880* and *CG13002* and downstream by RNA polymerase III subunit I *(Polr3I)* and Nitrogen permease regulator-like 2 *(Nprl2)*. The *tblastn* search of *D. melanogaster* DENR-PA (query) against the *D. simulans* (GenBank Accession: GCA_000754195.3) Genome Assembly (database) placed the putative ortholog of *DENR* within scaffold CM002914 (CM002914.1) at locus LOC6726200 (XP_039152449.1)— with an E-value of 5e-33 and a percent identity of 91.43%. Furthermore, the putative ortholog is flanked upstream by LOC6726201 (XP_002107176.1) and LOC6726203 (XP_002107178.3), which correspond to *CG4880* and *CG13002* in *D. melanogaster* (E-value: 0.0 and 1e-147; identity: 87.39% and 90.57%, respectively, as determined by *blastp*; **Figure 1A**, Altschul et al., 1990). The putative ortholog of *DENR* is flanked downstream by LOC6726199 (XP_002107174.1) and LOC6726198 (XP_002107173.1), which correspond to *Polr3I* and *Nprl2* in *D. melanogaster* (E-value: 8e-150 and 0.0; identity: 90.04% and 99.76%, respectively, as determined by *blastp*). The putative ortholog assignment for *DENR* in *D. simulans* is supported by the following evidence: the synteny of the genomic neighborhood is completely conserved across both species, and all *BLAST* search results used to determine orthology indicate very high-quality matches.

**Figure 1:**
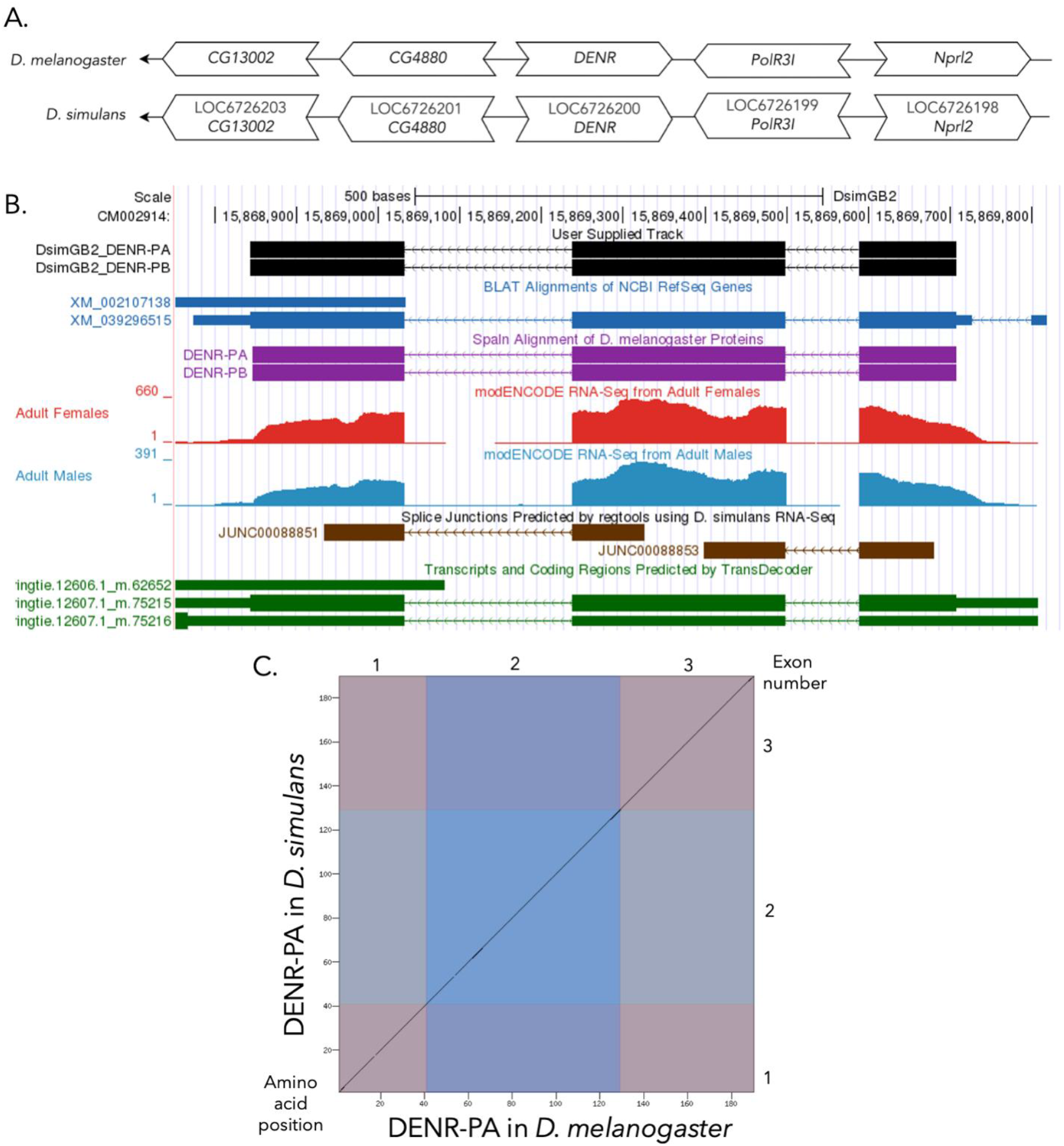
(A) Synteny comparison of the genomic neighborhoods for *DENR* in *Drosophila melanogaster* and *D. simulans*. Thin underlying arrows indicate the DNA strand within which the target gene–*DENR*–is located in *D. melanogaster* (top) and *D. simulans* (bottom). The thin arrows pointing to the left indicate that *DENR* is on the negative (-) strand in *D. melanogaster* and *D. simulans*. The wide gene arrows pointing in the same direction as *DENR* are on the same strand relative to the thin underlying arrows, while wide gene arrows pointing in the opposite direction of *DENR* are on the opposite strand relative to the thin underlying arrows. White gene arrows in *D. simulans* indicate orthology to the corresponding gene in *D. melanogaster*. Gene symbols given in the *D. simulans* gene arrows indicate the orthologous gene in *D. melanogaster*, while the locus identifiers are specific to *D. simulans*. **(B) Gene Model in GEP UCSC Track Data Hub** (Raney et al. 2014). The coding-regions of *DENR* in *D. simulans* are displayed in the User Supplied Track (black); coding exons are depicted by thick rectangles and introns by thin lines with arrows indicating the direction of transcription. Subsequent evidence tracks include BLAT Alignments of NCBI RefSeq Genes (dark blue, alignment of Ref-Seq genes for *D. simulans*), Spaln of D. melanogaster Proteins (purple, alignment of Ref-Seq proteins from *D. melanogaster*), Transcripts and Coding Regions Predicted by TransDecoder (dark green), RNA-Seq from Adult Females and Adult Males (red and light blue, respectively; alignment of Illumina RNA-Seq reads from *D. simulans*), and Splice Junctions Predicted by regtools using *D. simulans* RNA-Seq (Graveley et al. 2011; SRP006203). Splice junctions shown have a minimum read-depth of 10 with 500-999 supporting reads indicated in brown. **(C) Dot Plot of DENR-PA in *D. melanogaster* (*x*-axis) vs. the orthologous peptide in *D. simulans* (*y*-axis)**. Amino acid number is indicated along the left and bottom; coding-exon number is indicated along the top and right, and exons are also highlighted with alternating colors.

### Protein Model

*DENR* in *D. simulans* has two protein-coding isoforms (DENR-PA and DENR-PB; **Figure 1B**). Isoforms DENR-PA and DENR-PB are identical and contain three protein-coding exons. Relative to the ortholog in *D. melanogaster*, the coding-exon number is conserved, as DENR-PA and DENR-PB are also identical with three coding exons in *D. melanogaster*. The sequence of DENR-PA in *D. simulans* has 98.94% identity (E-value: 5e-138) with the protein-coding isoform DENR-PA in *D. melanogaster*, as determined by *blastp* (**Figure 1C**).

## Methods

“Detailed methods including algorithms, database versions, and citations for the complete annotation process can be found in Rele et al. (2023). Briefly, students use the GEP instance of the UCSC Genome Browser v.435 (https://gander.wustl.edu; Kent WJ et al., 2002; Navarro Gonzalez et al., 2021) to examine the genomic neighborhood of their reference IIS gene in the *D. melanogaster* genome assembly. Students then retrieve the protein sequence for the *D. melanogaster* reference gene for a given isoform and run it using *tblastn* against their target *Drosophila* species genome assembly on the NCBI BLAST server (https://blast.ncbi.nlm.nih.gov/Blast.cgi; Altschul et al., 1990) to identify potential orthologs. To validate the potential ortholog, students compare the local genomic neighborhood of their potential ortholog with the genomic neighborhood of their reference gene in *D. melanogaster*. This local synteny analysis includes at minimum the two upstream and downstream genes relative to their putative ortholog. They also explore other sets of genomic evidence using multiple alignment tracks in the Genome Browser, including BLAT alignments of RefSeq Genes, Spaln alignment of *D. melanogaster* proteins, multiple gene prediction tracks (e.g., GeMoMa, Geneid, Augustus), and modENCODE RNA-Seq from the target species. Detailed explanation of how these lines of genomic evidence are leveraged by students in gene model development is described in Rele et al. (2023). Genomic structure information (e.g., CDSs, intron-exon number and boundaries, number of isoforms) for the *D. melanogaster* reference gene is retrieved through the Gene Record Finder (https://gander.wustl.edu/~wilson/dmelgenerecord/index.html; Rele et al., 2023). Approximate splice sites within the target gene are determined using *tblastn* using the CDSs from the *D. melanogaste*r reference gene. Coordinates of CDSs are then refined by examining aligned modENCODE RNA-Seq data, and by applying paradigms of molecular biology such as identifying canonical splice site sequences and ensuring the maintenance of an open reading frame across hypothesized splice sites. Students then confirm the biological validity of their target gene model using the Gene Model Checker (https://gander.wustl.edu/~wilson/dmelgenerecord/index.html; Rele et al., 2023), which compares the structure and translated sequence from their hypothesized target gene model against the *D. melanogaster* reference gene model. At least two independent models for a gene are generated by students under mentorship of their faculty course instructors. Those models are then reconciled by a third independent researcher mentored by the project leaders to produce the final model. Note: comparison of 5’ and 3’ UTR sequence information is not included in this GEP CURE protocol.” (Gruys et al., 2025 - micropub)

## Supporting information

Supplemental Files with FASTA, PEP, GFF

## Acknowledgements

We would like to thank Wilson Leung for developing and maintaining the technological infrastructure that was used to create this gene model and Laura K. Reed for overseeing the project.

## Supplemental Files

1. Zip file containing a FASTA, PEP, GFF files for the gene model

## Metadata

Bioinformatics, Genomics, *Drosophila*, Genotype Data, New Finding

